# Effects of parity, blood progesterone, and non-steroidal anti-inflammatory treatment on the dynamics of the uterine microbiota of healthy postpartum dairy cows

**DOI:** 10.1101/2020.05.18.101766

**Authors:** O. Bogado Pascottini, J. F. W. Spricigo, S. J. Van Schyndel, B. Mion, J. Rousseau, J. S. Weese, S. J. LeBlanc

## Abstract

This study evaluated the effects of treatment with meloxicam (a non-steroidal anti-inflammatory drug), parity, and blood progesterone concentration on the dynamics of the uterine microbiome of clinically healthy postpartum dairy cows. Seven primiparous and 9 multiparous postpartum Holstein cows received meloxicam (0.5 mg/kg SC, n = 7 cows) once daily for 4 days (10 to 13 days in milk (DIM)) or were untreated (n = 9 cows). Endometrial cytology samples were collected by cytobrush at 10, 21, and 35 DIM, from which the metagenomic analysis was done using 16S rRNA gene sequence analysis. A radioimmunoassay was used to measure progesterone concentration in blood serum samples at 35 DIM and cows were classified as > 1 ng/mL (n = 10) or ≤ 1 ng/mL (n = 6). Alpha diversity for bacterial genera (Chao1, Shannon-Weiner, and Camargo’s evenness indices) were not affected by DIM, meloxicam treatment, parity, or progesterone category (*P* > 0.2). For beta diversity (genera level), principal coordinate analysis (Bray-Curtis) showed differences in microbiome between parity groups (*P* = 0.01).

There was lower overall abundance of *Anaerococcus, Bifidobacterium, Corynebacterium, Lactobacillus, Paracoccus, Staphylococcus*, and *Streptococcus* and higher abundance of *Bacillus, Fusobacterium*, and *Novosphingobium* in primiparous than multiparous cows (*P* < 0.05); these patterns were consistent across sampling days. Bray-Curtis dissimilarity did not differ by DIM at sampling, meloxicam treatment, or progesterone category at 35 DIM (*P* > 0.5). In conclusion, uterine bacterial composition was not different at 10, 21, or 35 DIM, and meloxicam treatment or progesterone category did not affect uterine microbiota in clinically healthy postpartum dairy cows. Primiparous cows presented a different composition of uterine bacteria than multiparous cows. The differences in microbiome associated with parity might be attributable to changes that occur consequent to the first calving, but this hypothesis should be investigated further.

## Introduction

After parturition, the endometrial epithelium and caruncles are sloughed and the uterus of essentially all cows is exposed to potentially pathogenic bacteria [1, 2]. Up to half of dairy cows experience at least one form of reproductive tract inflammatory disease within 35 days in milk (DIM) [3]. This takes the form of systemic illness with fetid uterine discharge before 14 DIM (metritis), or localized infection of the uterus and/or cervix (diagnosed as purulent vaginal discharge (PVD) around 5 weeks postpartum), or localized inflammation diagnosed by cytology (without PVD) referred to as subclinical endometritis (SCE). The two former conditions are clearly associated with bacterial pathogens: metritis predominantly with Gram-negative anaerobes (e.g., *Fusobacterium* and *Porphorymonas*) [4], and PVD with *Trueperella pyogenes* [5]. SCE might be initiated by bacterial infection, but it is not consistently associated with that the uterine microbiome [5]. The point at which uterine inflammation and uterine microbiota shifts from physiologic to pathologic and the determinants of these processes are only partially understood [6]. Questions remain about which bacteria may be critical in pathogenesis, and how the interplay between bacterial infection and host response lead to or avoid disease [2].

Studies using conventional bacteriology methods rely on the ability to culture live bacteria that grow under controlled conditions in the lab, whereas genetic sequencing technologies allow identification of a fuller range of uterine bacteria [7]. The advent of culture-independent methods such as 16S ribosomal RNA gene sequencing has advanced knowledge about bacterial communities in the reproductive tract of cows [8]. Bacteria are found in the uterus of healthy cows, virgin heifers, and even in pregnant cows [9, 10]. Understanding the dynamics of the uterine microbiota and the factors that affect the microbiome may provide deeper insights as to the pathophysiology of uterine disease. However, it is currently difficult to define a “healthy” uterine microbiota [11]. In this regard, many physiological conditions such as resident uterine microbiota, time relative to calving, degree of metabolic stress, and concentrations of sex hormones or other changes inherent in the estrus cycle may play a role in the uterine bacterial composition of healthy postpartum cows.

In a previous study, we showed that treatment with the non-steroidal anti-inflammatory drug (NSAID) meloxicam (MEL) from 10 to 13 DIM transiently reduced haptoglobin (a marker of systemic inflammation) and improved indicators of energy metabolism (greater insulin-like growth factor-1 and glucose, and lesser β-hydroxybutyrate), and modestly increased circulating polymorphonuclear (PMN) cell function [12]. For the present analysis using samples from a subset of the same cows, the hypothesis was that metabolic changes induced by MEL would affect the uterine microbiota. Thus, our objective was to evaluate the effects of MEL on the uterine microbiota of clinically healthy postpartum dairy cows. We also aimed to study the effects of DIM at sampling, parity, and blood progesterone concentration on the dynamics of the uterine microbiome at 10, 21, and 35 DIM.

## Materials and methods

### Ethics approval

Animal handling procedures, sampling, and treatment were approved by the University of Guelph Animal Care Committee. Cows were managed according to the code of practice of the National Farm Animal Care Council.

### Study design

This study derives from a larger experiment on postpartum NSAID treatment [12] that was conducted from April to August 2018 at the University of Guelph Livestock Research and Innovation Centre, Dairy Facility (Elora, ON, Canada). For the larger experiment, 42 Holstein cows were used, but for this study, a subset of 16 Holstein cows was deliberately selected to balance for parity (n = 7, primiparous (PRIM) and n = 9, multiparous (MULT)), treatment (n = 7, MEL and n = 9, control (CON)), and progesterone serum concentration at 35 d in milk (DIM; n = 10, > 1 ng/mL (HIGH; functional corpus luteum present) and n = 6, ≤ 1 ng/mL (LOW; corpus luteum absent)).

### Management

Cows calved in individual box stalls and were moved to free-stall pens 5 days after calving. Cows were fed *ad libitum* from individually assigned automated feed bins (Insentec B.V., Marknesse, the Netherlands), and were milked twice daily in a rotary parlor. Only cows considered clinically healthy from calving to 35 DIM were included (unassisted calving and absence of retained placenta, metritis, or other clinical disease (including PVD) before or during the study period). Treated cows received subcutaneous injections of MEL (0.5 mg/kg of body weight; Metacam, Boehringer Ingelheim Canada Ltd., Burlington, ON, Canada) once daily at 10, 11, 12, and 13 DIM. CON cows did not receive any injections.

### Blood sampling and serum progesterone analysis

Blood samples were collected by coccygeal venipuncture into vacuum tubes without anticoagulant (BD Vacutainer Precision Glide, Becton Dickinson, Franklin Lakes, NJ) at 35 DIM. All blood samples were allowed to clot, centrifuged at 1,500 × g for 15 min and serum was stored in aliquots at -20°C. Progesterone was measured using the Progesterone Double Antibody RIA Kit (MPBiomedicals, Inc., Costa Mesa, CA, USA) according to the manufacturer’s instructions. Briefly, 100 μL of each standard, serum sample, and control were added into their respective anti-progesterone coated tubes. Then, 1 mL of progesterone I-125 was added to all tubes and vortexed briefly. Tubes were then incubated in a 37°C-water bath for 120 min. The contents of the tubes were decanted, and tubes were counted in a gamma counter calibrated for I-125 (Biodex Medical Systems, Inc.). The lower limit of detection was 0.1 ng/ml, and the intra-assay coefficient of variation was 14.3%. Samples were analyzed in the same batch.

### Cytology collection

Endometrial cytobrush samples were collected at 10, 21, and 35 DIM. Briefly, after cleansing the perineal area of the cow with antibacterial soap and water, 70% isopropyl alcohol was sprayed and dried thoroughly with paper towels. A sterile cytobrush rod (covered by a sterile sanitary sheath) was introduced into the vagina and guided through the cervix via rectal palpation. Once the tip of the rod reached the uterine body, the sanitary sheath was pulled back, the cytobrush was exposed from the rod and rotated against the dorsal wall of the endometrium (*corpus uteri* region) with gentle pressure from the index finger through the rectum. The cytobrush was retracted into the rod and removed from the vagina. Once outside the genital tract, the cytobrush was gently rolled onto a sterilized microscope slide. The cytobrush tip was then cut with sterilized scissors, inserted into a sterile 2 mL cryovial, and stored at -80°C within 15 min. After cytobrush sampling, vaginal discharge was evaluated using the Metricheck device (Simcrotech, Hamilton, New Zealand) and scored as 0 = clear mucus, 1 = mucus with flecks of pus, 2 = mucopurulent discharge (≤ 50% pus), and 3 = purulent discharge (> 50% pus). Cytobrush microscope slides were stained using May-Grunwald-Giemsa stain, 300 cells were counted per slide in multiple fields, and the proportion of PMN cells to epithelial cells (PMN%) was determined.

### DNA extraction

DNA was extracted using the QIAamp Microbiome kit (Qiagen Inc.; Toronto, ON, Canada) following the instructions of the manufacturer. DNA concentration was measured by a NanoDrop ND-1000 uv-vis spectrophotometer. (Thermo Fisher Scientific; Wilmington DE, USA) based on the absorbance at 260 nm and using the Beer-Lambert equation. The DNA quality was assessed by using the 260/280 nm ratio.

### 16S rRNA Gene Amplification and Sequencing

PCR amplification of the V4 hypervariable region of the 16S rRNA gene was performed using the forward primer 515F-mod and reverse primer 806R-mod [13]. The forward and reverse primers were designed to contain an Illumina® overhang adapter sequence (Illumina®; San Diego, CA, USA) in order to anneal them to primers containing the Illumina® adaptors plus the 8 bp identifier indices. The following PCR conditions were used: 3 minutes at 94°C for denaturing, followed by 30 cycles of 45 s at 94°C, 60 s at 52°C, and 60 s at 72°C, with a final elongation of 10 minutes at 72°C. After amplification, the PCR products were evaluated by electrophoresis in 1.5% agarose gel and purified with the Mag-Bind RxnPure Plus kit (Omega Bio-Tek, CA, Norcross, GA, USA) by mixing 20 μL of amplicon with 25 μL of Mag-Bind in a 96 well flat-bottom micro-titer plate. After incubation for 5 min at room temperature, the beads were separated and washed twice with 80% ethanol and then eluted in 32 μL of 10 mM Tris pH 8.5 buffer solution. A second PCR was performed to attach dual indices and Illumina® sequencing adapters using the Nextera XT Index kit (Illumina®; San Diego, CA, USA). The conditions of this PCR included: 3 minutes at 94°C followed by 8 cycles of 30 s at 94°C, 30 s at 55°C, and 30 s at 72°C, plus a final step of 10 minutes at 72°C. After purification of these amplicons, the samples were quantified via NanoDrop® spectrophotometry (Thermo Fisher Scientific; Wilmington DE, USA). Normalization and sequencing of the library pool was performed at the Agriculture and Food Laboratory, University of Guelph using an Illumina® MiSeq Reagent kit with a 2 × 250 bp read length.

### Statistical analysis

The statistical analyses were performed using RStudio (version 3.6.3; R Core Team, Vienna, Austria) by employing the packages vegan, fossil, and phyloseq. First, decontamination was done via the decontam R package by using the sequencing of 6 sterile cytobrush samples exposed to the farm environment (Supplemental Figure S1). Alpha diversity for bacterial genera (Chao1, Shannon-Weiner, and Camargo’s evenness indices) was compared by DIM at sampling (10 vs. 21 vs. 35), meloxicam treatment (CON vs. MEL), parity (PRIM vs. MULT), and serum progesterone category at 35 DIM (LOW vs. HIGH) using Kruskal–Wallis analysis of variance. For beta diversity (phylum and genera levels), principal coordinate analysis (Bray-Curtis) was used to assess differences in uterine bacterial composition by DIM, using non-parametric multivariate analysis of variance (PERMANOVA). The interaction of DIM at sampling with MEL, parity, and serum progesterone category at 35 DIM was evaluated with two-way PERMANOVA with 1000 permutations based on Bray-Curtis dissimilarity. Venn diagrams were generated to show the number of bacterial core genera (having > 1% abundance) by DIM at sampling and by MEL, parity, and serum progesterone category at 35 DIM. The *simper* function (vegan package in R) was used to set the similarity percentages to extract the most influential bacteria (phylum and genera levels) and compare their relative abundance by DIM at sampling and MEL, parity, and serum progesterone category at 35 DIM (Kruskal–Wallis analysis of variance). Heatmaps representing bacteria phyla and genera fold-changes (average linkage clustering based on Bray–Curtis distance) and similarity dendrograms (based on Bray– Curtis distance and unweighted pair group method with arithmetic mean clustering) were built to show bacterial clustering (package Heatplus in R).

## Results

### Descriptive statistics

The data for this non-randomized clinical trial were from 16 healthy Holstein cows that completed the transition period (from calving until 35 DIM) without clinical disease or any antibiotic treatment. The detailed composition of the experimental groups including prepartum BCS, BCS at 35 DIM, average milk production to 35 DIM, average parity per experimental group, and average progesterone concentration at 35 DIM experimental group are shown in Supplemental Table S1. None of the cows had PVD at 35 DIM (mucopurulent discharge or worse) and the proportion of endometrial PMN at 10, 21, and 35 DIM per experimental group is summarized in Supplemental Table S2.

### DIM at sampling

Figure 1A shows no differences in alpha diversity indices (Chao1, Shannon-Weiner, and Camargo’s evenness indices) for bacteria genera at 10, 21, and 35 DIM. The top 10 bacteria relative abundance and the number of bacterial core genera were similar among samples taken at 10, 21, and 35 DIM (Figures 1B and 1C, respectively). The relative abundance of the most influential bacteria phyla was not affected by DIM at sampling (Supplemental Figure S2). Principal coordinate analysis based on Bray-Curtis dissimilarity (beta diversity) shows a similar microbiome composition across sampling days at the genera (Figure 1) and phyla levels (Supplemental Figure S3).

**Figure 1.**
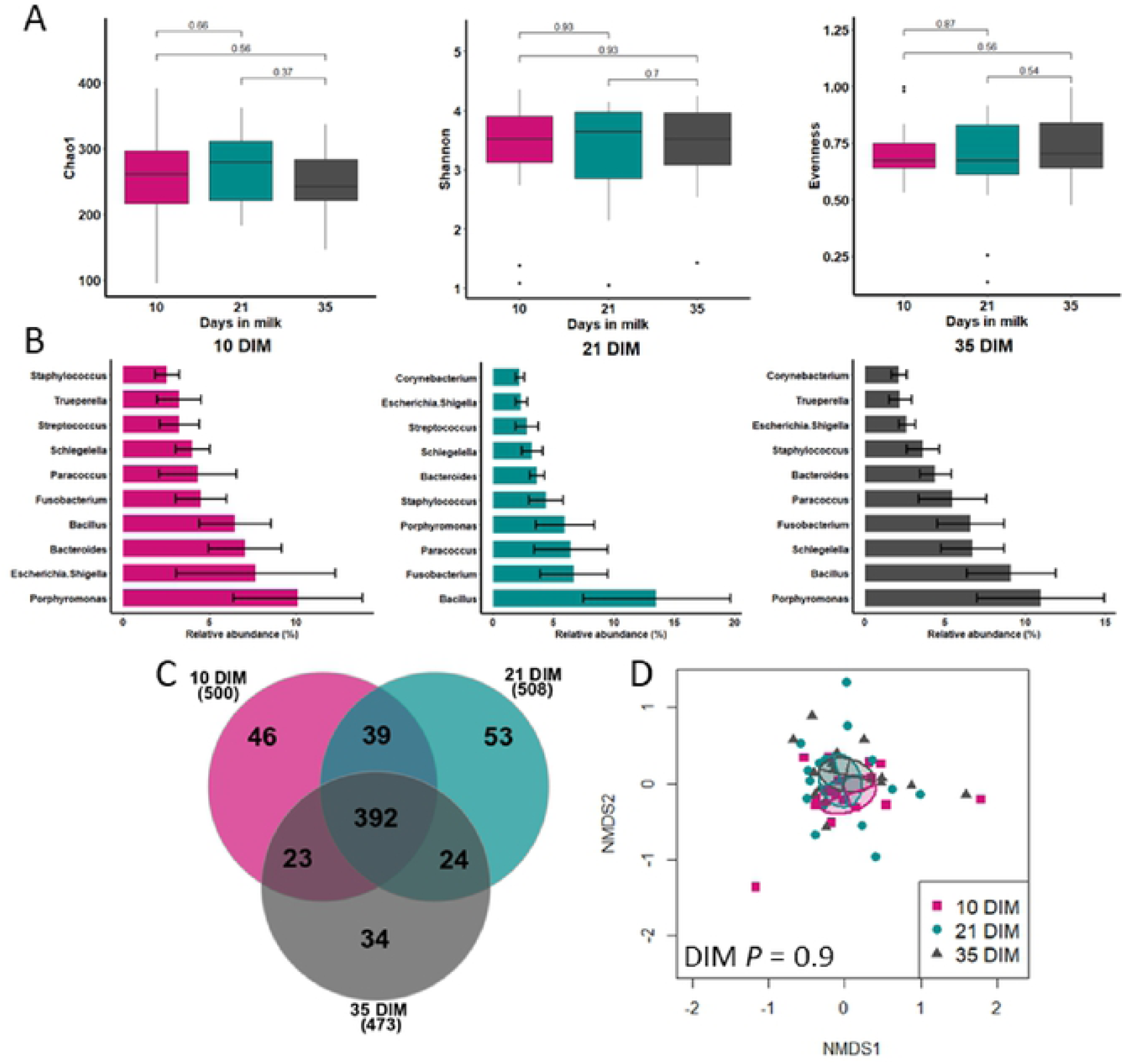
Dynamics of uterine microbiota in clinically healthy postpartum dairy cows (n = 16) in samples collected at 10, 21, and 35 d in milk (DIM). A) Alpha diversity for bacterial genera (numbers above the brackets are *P* values), B) mean relative abundance (with standard error) of the top ten bacteria genera, C) number of bacterial core genera (having> 1% abundance), and D) beta diversity (principal coordinate analysis (Bray-Curtis)) were similar among 10, 21, and 35 DIM.

**Figure 2.**
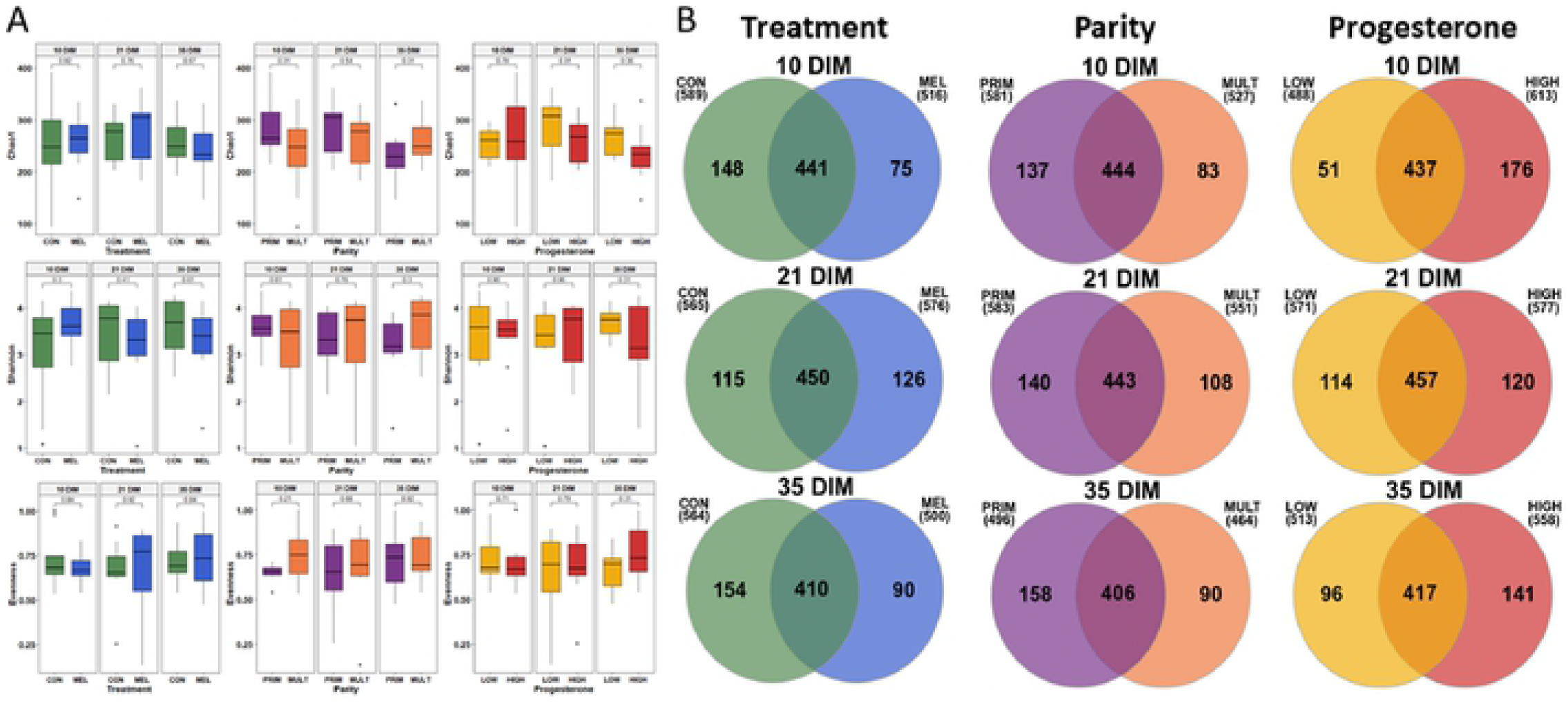
Effects of meloxicam treatment (n = 9, control (CON) and n = 7, meloxicam (MEL)), parity (n = 7, primiparous (PRIM) and n = 9, multiparous (MULT)), and blood progesterone concentration at 35 DIM (n = 10,> 1 ng/mL (HIGH) and n = 6, ≤ 1 ng/mL (LOW)) on the dynamics of the uterine microbiota of clinically healthy postpartum dairy cows (n **=** 18) in samples collected at 10, 21, and 35 d in milk (DIM). A) Alpha diversity for bacterial genera was not affected by MEL, parity, or progesterone category at 35 DIM (numbers above the brackets are *P* values). B) There number of bacterial core genera (having> 1% abundance) was variable between CON and MEL. PRIM and LOW had more stable, greater numbers of bacterial core genera than MULT and HIGH, respectively.

### MEL, parity, and blood progesterone

Alpha diversity for bacterial genera was not associated with MEL, parity, or progesterone category at 35 DIM (Figure 3A). The number of bacterial core genera (Venn diagram) stratified by MEL, parity, and progesterone category at 35 DIM are illustrated in Figure 3B. Principal coordinate analysis for Bray-Curtis dissimilarity (genus level) was not affected by MEL or progesterone category at 35 DIM (Figure 3). Genera beta diversity (Bray-Curtis dissimilarity) showed differences in microbiome composition between parity groups (Figure 3). Beta diversity at the phyla level was not affected by MEL, parity, or progesterone category at 35 DIM (Supplemental Figure S4). The relative abundance of the most influential bacteria phyla and genera are shown in Figures 4 and 5 and in Supplementary Figures S5 to S10. MULT had greater abundance of *Actinobacteria* than PRIM (Supplementary Figures S6 and S8). There was lower overall abundance of *Anaerococcus, Bifidobacterium, Corynebacterium, Lactobacillus, Paracoccus, Staphylococcus*, and *Streptococcus* and higher abundance of *Bacillus, Fusobacterium*, and *Novosphingobium* in PRIM than MULT (Figures 4 and 5). The heatmap and similarity dendrograms in Figure 6 show a more similar clustering for bacterial genera in parity groups than for MEL or progesterone category at 35 DIM. At the phyla level (Supplemental Figure S11), heatmap and similarity dendrograms show similar tendencies as for the genera level (more similar clustering in parity groups).

**Figure 3.**
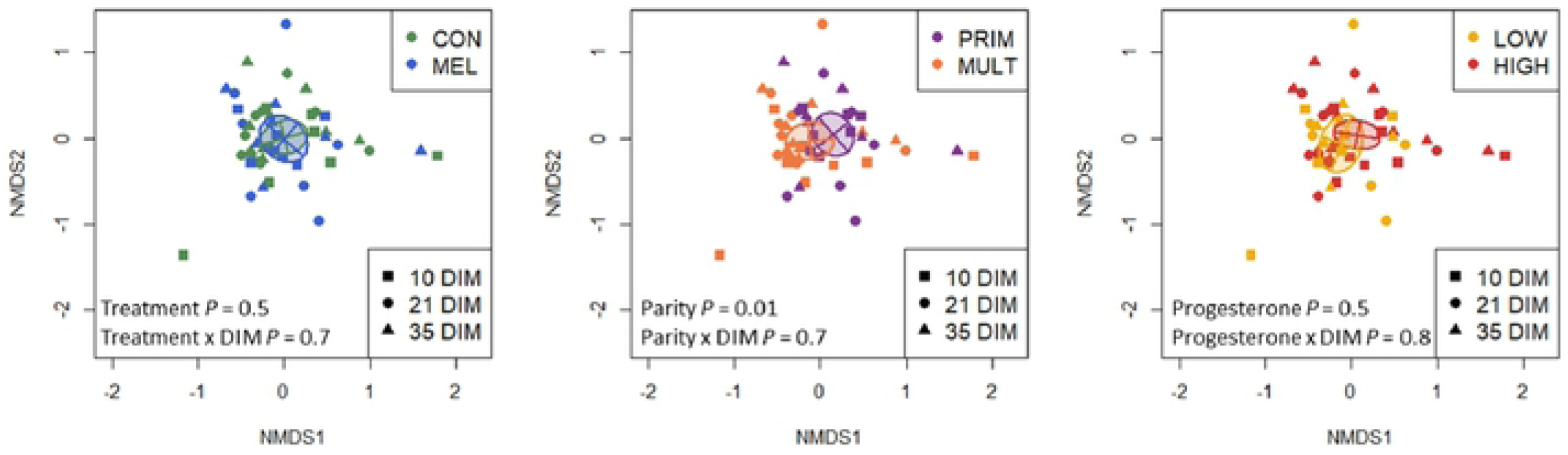
Effects of meloxicam treatment (n = 9, control (CON) and n = 7, meloxicam (MEL)), parity (n = 7, primiparous (PRIM) and n = 9, multiparous (MULT)), and blood progesterone concentration at 35 DIM (n = 10, > 1 ng/mL (HIGH) and n = 6, ≤ 1 ng/mL (LOW)) on the dynamics of the uterine microbiota of clinically healthy postpartum dairy cows (n = 18) in samples collected at 10, 21, and 35 d in milk (DIM). Principal coordinate analysis for Bray-Curtis dissimilarity (genus level) was not affected by MEL or progesterone concentration at 35 DIM. Genera beta diversity (Bray-Curtis dissimilarity) showed differences in microbiome composition between parity groups.

**Figure 4.**
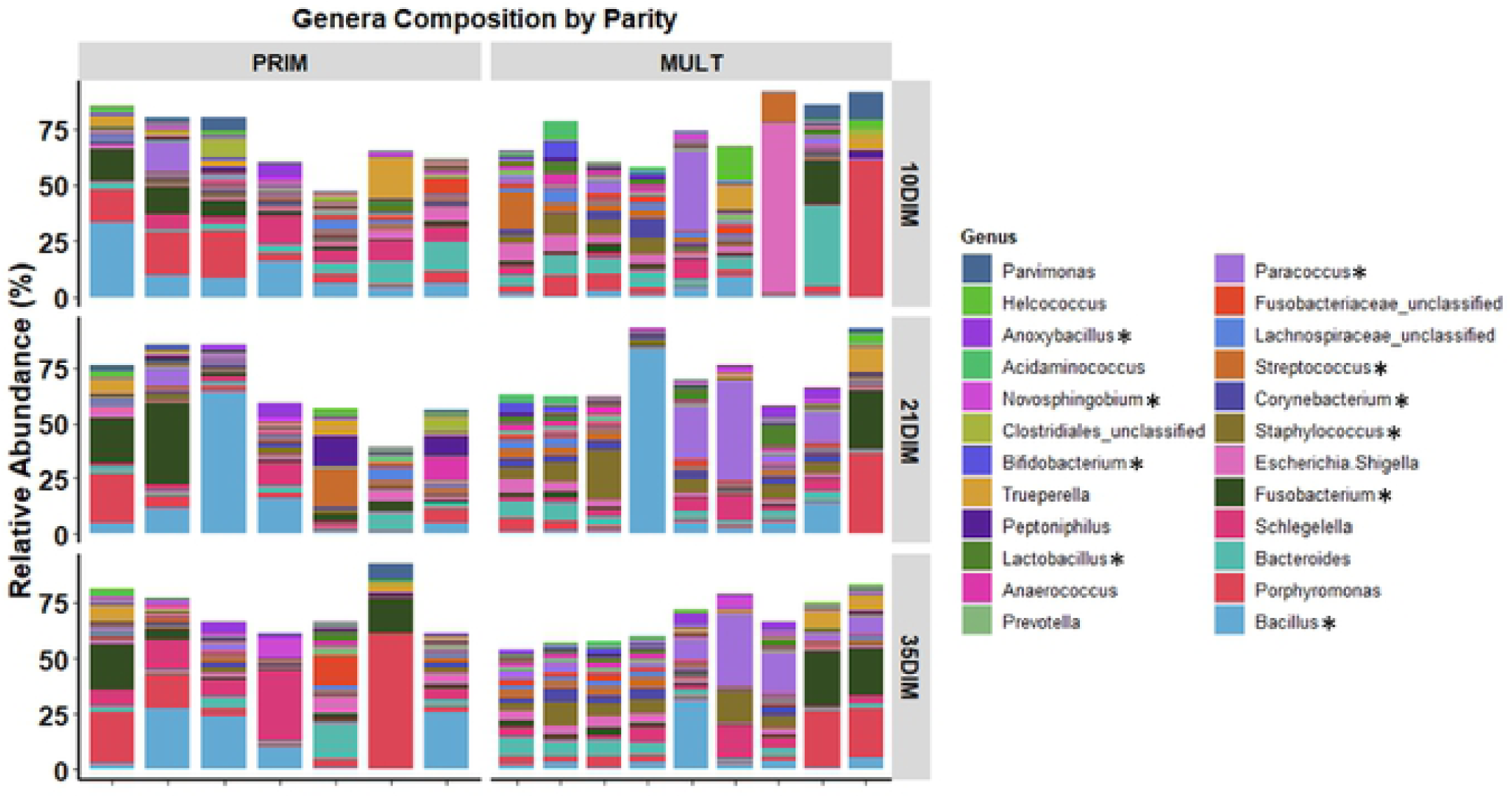
Relative abundance of the most influential bacterial genera in clinically healthy primiparous (PRIM, n = 7) and multiparous (MULT, n = 9) postpartum dairy cows in uterine samples collected at 10, 21, and 35 d in milk (DIM). There was an overall lower abundance of *Anaerococcus, Bifidobacterium, Corynebacterium, Lactobacillus, Paracoccus, Staphylococcus*, and *Streptococcus* and higher abundance of *Bacillus, Fusobacterium*, and *Novosphingobium* in PRIM than MULT *(P*< 0.05). Asterisks indicate differences between parity groups.

**Figure 5.**
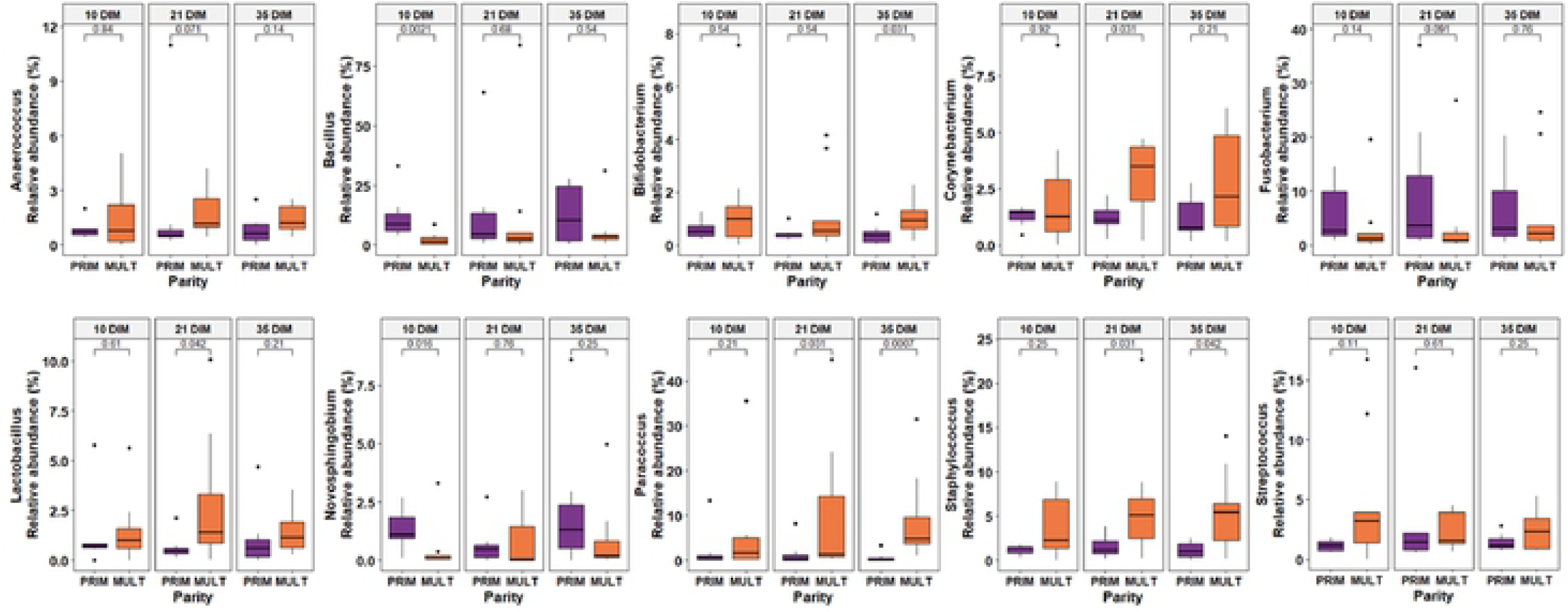
Relative abundance (by days in milk (DIM)) of bacterial genera that differed between clinically healthy primiparous (PRIM, n = 7) and multiparous (MULT, n = 9) postpartum dairy cows in uterine samples collected at 10, 21, and 35 DIM (numbers above the brackets are *P* values).

**Figure 6.**
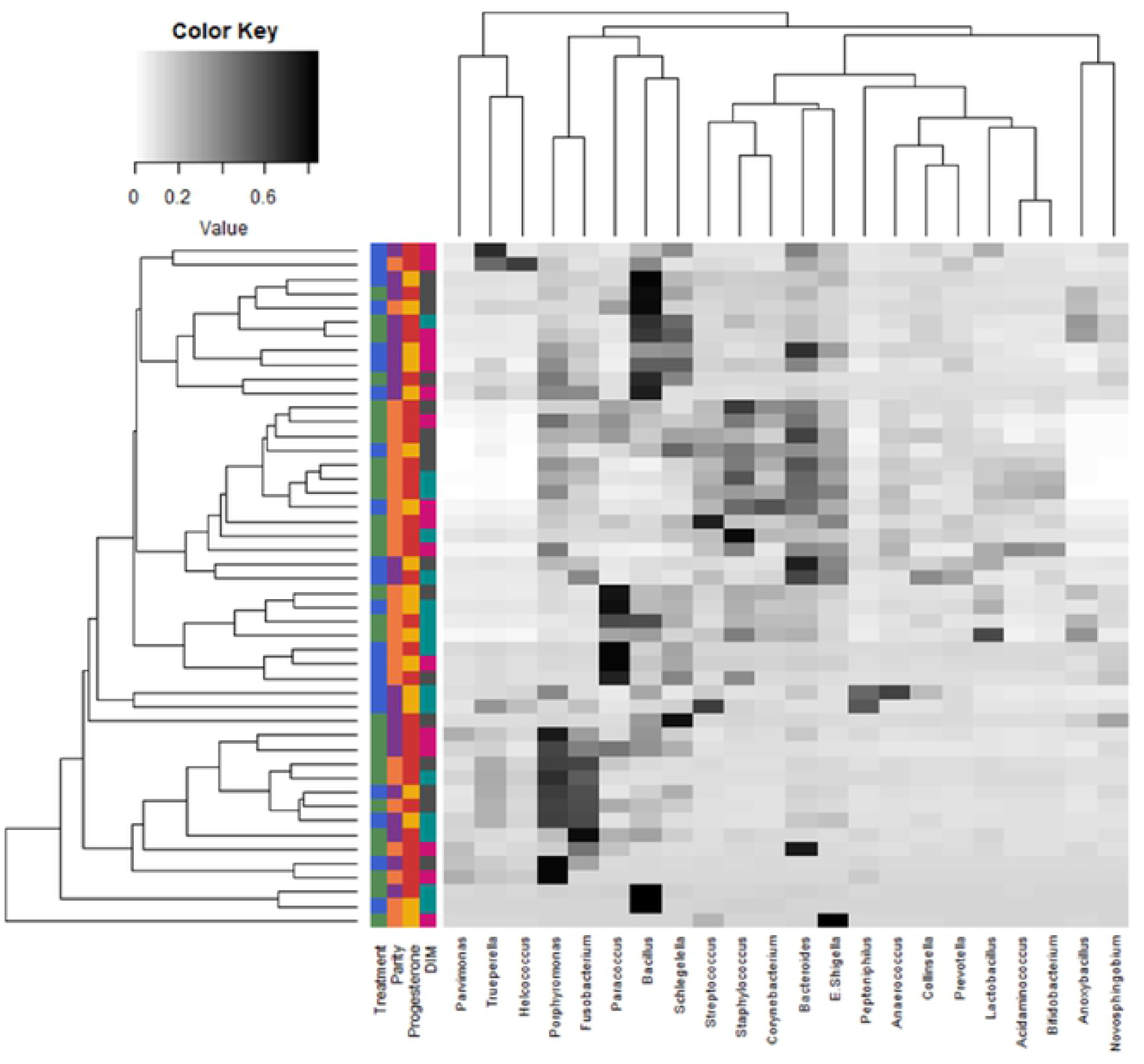
Heatmap with average linkage clustering based on Bray-Curtis distance showing the relative abundances of the most influential uterine bacteria genera in clinically healthy postpartum dairy cows (n = 18) in samples collected at 10, 21, and 35 d in milk (DIM). The relative abundance is indicated by a gradient of color from black (high abundance) to white (low abundance). Similarity dendrogram shows the unweighted pair group method with arithmetic mean (UPGMA) clustering of bacterial genera by treatment (n = 9, control (green) and n = 7, meloxicam (blue)), parity (n = 7, primiparous (purple) and n = 9, multiparous (orange), blood progesterone concentration at 35 DIM (n = 10, > 1 ng/mL (red) and n = 6, ≤ 1 ng/mL (yellow)), and DIM (10 DIM (pink), 21 DIM (turquoise), and 35 DIM (dark grey)).

## Discussion

This study focused on describing the effects of NSAID treatment and other factors that may affect the dynamics of bacterial populations by using 16S rRNA pyrosequencing of endometrial samples obtained from the postpartum uterus of clinically healthy dairy cows. Neither MEL (from 10 to 13 DIM) nor DIM at sampling (10, 21, and 35) influenced the structure or composition of the microbiome found in the postpartum uterus. Primiparity was associated with a more diverse composition of uterine bacteria than multiparity, with differences in the abundance of multiple bacterial genera. Progesterone category at 35 DIM did not have an effect on the uterine microbiome.

We aimed to investigate the uterine bacteria dynamics in clinically healthy dairy cows after the vaginal-uterine decompartmentalization process. In this regard, Miranda-CasoLuengo et al. (2019) [14] demonstrated that compartmentalization between the vagina and the uterine microbiome is re-established after 7 DIM in clinically healthy cows. Therefore, we decided to analyze the microbiota of our uterine samples starting at 10 DIM. Sampling the endometrium within the first week postpartum can be challenging. Furthermore, based on our experience from previous experiments, uterine sampling within the first week postpartum can be difficult because in early involution many cows have large, pendulous uteri that cannot be properly catheterized.

In humans, transcervical uterine sampling is described as a cause of cross-contamination with the cervical microbiota [11]. Excessive manipulation of the genital tract should be avoided, and cervical catheterization should be done efficiently and without force.

Certain generalizations can be made from the existing literature, but there is no consensus on the nature of a “healthy” uterine microbiota [11]. Although none of the cows in our study had clinical disease, there was variation in the proportions of endometrial PMN. We attempted to balance the parity and progesterone groups for endometrial PMN proportions. Importantly, Wang et al. (2018) [5] showed that abundances of common bacteria or putative pathogens were not associated with high proportions of PMN (> 18%) at 30 DIM using 16S rRNA pyrosequencing. On the other hand, it was shown that in cows with PVD, there was an increased relative abundance of *Trueperella* in uterine samples collected between 25 and 35 DIM [5, 15]. None of the cows in this study had PVD (mucopurulent vaginal discharge or worse) at 35 DIM. Therefore, we consider that our results are not confounded by other factors than our variables of interest.

Our results clearly show that a uterine microbiota was already established at 10 DIM, and its composition did not change at 21 or 35 DIM. At each time point, *Firmicutes* was the most abundant phylum followed by *Proteobacteria, Bacteroidetes*, and *Actinobacteria*. At the genus level, *Bacillus, Fusobacterium*, and *Porphyromonas* were (interchangeably) within top 5 bacteria with the greatest relative abundance. However, the sum of their relative abundances was less than 30% of the total bacteria population. This highlights the tremendous bacterial diversity encountered by the uterus of clinically healthy postpartum cows. Increased abundance of *Bacteroides, Porphyromonas*, and *Fusobacterium* accompanied with loss of heterogeneity was found in cows that later developed metritis [8]. The total relative abundance of these bacterial genera was over 60% of the uterine microbiota of metritic cows [9]. Potential pathogenic bacteria are present in the uterus of postpartum cows, but our results, along with others, indicate that high microbial diversity is the key to a healthy uterine environment.

The postpartum metabolic milieu of the cow likely contributes to development of distinct microbiota in the reproductive tract [14]. The samples for the microbiome analyses for this study were derived from a larger experiment where postpartum MEL reduced systemic inflammation and improved indicators of energy metabolism [12]. Meloxicam treatment slightly enhanced circulating PMN function, but the endometrial inflammatory status was not affected. Treatment with NSAID could affect the composition of the uterine microbiome, either directly or indirectly. Antibacterial properties of some NSAID were recently discovered [16]. Diclofenac, aspirin, and etodolac could prevent biofilm formation at normal doses used during anti-inflammatory therapy [17]. However, the mechanisms of antimicrobial activity of NSAID are mostly unclear.

Traditionally, the therapeutic use of NSAID is effective at blocking the conversion of arachidonic acid to prostaglandins by inhibiting the inducible cyclooxygenase-2 pathway [18]. Moreover, it was also demonstrated that NSAID could down-regulate the inflammatory response by inhibiting pro-inflammatory cytokine production, as well as suppression of nuclear factor-kappa B activity [19]. Therefore, if we reduced systemic and uterine inflammation, this would potentially indirectly affect uterine bacterial composition. However, MEL did not affect the endometrial inflammatory status. Consequently, here we show that MEL affected neither the alpha or beta microbiome diversities nor relative abundances of bacteria at the phylum or genera levels. Nevertheless, it is important to mention that our MEL treatment was from 10 to 13 DIM, so from the last MEL injection to the next endometrial sample for 16S rRNA pyrosequencing there was a lag of 7 days. Thus, this study demonstrated the lack of effect of MEL on uterine microbiota in the medium term. The short-term effect of NSAIDs on the uterine microbiota remains uncertain.

Our principal coordinate analysis demonstrates that the uterine microbiomes from PRIM and MULT differ from each other, with a more diverse composition in cows that recently calved for the first time. The differences in the microbiome associated with parity could have several explanations. In women, a distinct vaginal microbiome prior to conception influences the relative composition of the vaginal microbiota during the first trimester of pregnancy [20]. The difference between parities might be attributable to resident bacteria that have already been modified in cows that have previously calved. Also, differences in anatomy and involution between older cows and those just calved for the first time may affect the vaginal-uterine bacterial exchange. The differences in the microbiome associated with parity might be attributable to changes occurring at parturition. Bicalho et al. (2017) [21] found greater diversity but with lesser total bacterial counts in vaginal samples collected just after calving in MULT than PRIM. The increased likelihood of bacterial contamination in PRIM could be related to a longer or more difficult parturition (although eutocic), resulting in increased trauma or altered neutrophil function [22, 23]. On the other hand, differences in metabolism and the characteristic greater milk production in MULT could also contribute to differences in their uterine bacterial composition. MEL induced changes in the metabolic profile by decreasing haptoglobin and improving markers of energy metabolism, but with lack of effects on the uterine microbiota. However, these metabolic changes were transient and returned to basal levels just after the end of anti-inflammatory treatment. Metabolic adaptations associated with milk production associated with multiparity are more profound and persistent.

This study shows that progesterone serum category at 35 DIM was not associated with distinct uterine microbiota. We hypothesized that the endometrial changes associated with increased circulating progesterone concentration (indicating that ovulation had occurred) could affect the uterine microbiome [24]. Although few cows are expected to have a functional corpus luteum by 10 or 21 DIM, we did not measure the blood progesterone concentrations at these time points. Our data do not refute possible effects of progesterone on the uterine microbiome later in lactation in estrus cyclic cows.

## Conclusions

The uterine microbiota in endometrial cytobrush samples collected from clinically healthy postpartum dairy cows was not different at 10, 21, or 35 DIM. Meloxicam treatment from 10 to 13 DIM did not affect the uterine microbiota. There was greater diversity in PRIM than MULT, with greater overall abundance of *Bacillus, Fusobacterium*, and *Novosphingobium* and lesser abundance of *Anaerococcus, Bifidobacterium, Corynebacterium, Lactobacillus, Paracoccus, Staphylococcus*, and *Streptococcus*. Progesterone serum category at 35 DIM was not associated with the uterine bacterial composition. We encourage further characterization of the establishment of the uterine microbiome in young cattle, and how it changes with the first and subsequent calvings.

## Acknowledgments

Financial support for this study was provided by the Ontario Ministry of Agriculture, Food and Rural Affairs (Guelph, ON, Canada) and by Boehringer-Ingelheim Vetmedica (Burlington, ON, Canada). Special thanks to Laura Wright (University of Guelph, Guelph, ON, Canada) and the staff at the University of Guelph Livestock Research and Innovation Centre Dairy Facility (Elora, ON, Canada).

